# Maximizing statistical power to detect clinically associated cell states with scPOST

**DOI:** 10.1101/2020.11.23.390682

**Authors:** Nghia Millard, Ilya Korsunsky, Kathryn Weinand, Chamith Y. Fonseka, Aparna Nathan, Joyce B. Kang, Soumya Raychaudhuri

## Abstract

As advances in single-cell technologies enable the unbiased assay of thousands of cells simultaneously, human disease studies are able to identify clinically associated cell states using case-control study designs. These studies require precious clinical samples and costly technologies; therefore, it is critical to employ study design principles that maximize power to detect cell state frequency shifts between conditions, such as disease versus healthy. Here, we present single-cell Power Simulation Tool (scPOST), a method that enables users to estimate power under different study designs. To approximate the specific experimental and clinical scenarios being investigated, scPOST takes prototype (public or pilot) single-cell data as input and generates large numbers of single-cell datasets *in silico*. We use scPOST to perform power analyses on three independent single-cell datasets that span diverse experimental conditions: a batch-corrected 21-sample rheumatoid arthritis dataset (5,265 cells) from synovial tissue, a 259-sample tuberculosis progression dataset (496,517 memory T cells) from peripheral blood mononuclear cells (PBMCs), and a 30-sample ulcerative colitis dataset (235,229 cells) from intestinal biopsies. Over thousands of simulations, we consistently observe that power to detect frequency shifts in cell states is maximized by larger numbers of independent clinical samples, reduced batch effects, and smaller variation in a cell state’s frequency across samples.

## Introduction

Single-cell technologies are revolutionizing biological studies. For example, single-cell RNA sequencing (scRNA-seq)^1,2^ enables the measurement of the entire transcriptome of individual cells to reveal previously unobserved cell state heterogeneity^3,4^. Technological advances allowing for simultaneous measurement of an increasing diversity of modalities^5–8^ from increasing numbers of cells^9^ are enabling disease-focused single-cell studies to identify cell states that correlate with disease in blood or tissue^10–15^. Shifts in cell state frequencies (expansions or depletions) are often associated with biologically meaningful phenotypes; expanded populations in disease states are often associated with pathogenic mechanisms, which can be further defined by differential gene expression programs^16^. For example, a recent study identified and characterized a CD14^+^ monocyte state expanded in individuals with sepsis versus healthy controls^11^. Furthermore, differentially abundant cell states have also been associated with genetic variants^17^. In order to efficiently detect disease-associated cell states, it is essential to conduct large, well-powered case-control comparisons^18^.

In this study, we consider a wide range of factors that potentially affect power: variation in cell state frequencies across samples, covariation and inter-sample variation in gene expression, batch variability and structure, number of cells and samples, and sequencing depth. Due to variation across studies, these factors must be simulated in the study’s context to accurately estimate power.

Current single-cell simulation strategies directly simulate individual genes and focus on estimating power to detect general differential gene expression^19–23^ or associations with genetic variants^23–24^. These strategies are typically used to simulate small datasets ranging from 400-10,000 cells with few replicates, because sampling individual genes is slow and limits larger simulations. Furthermore, no strategies model the inter-sample variation in cell state frequencies across samples that is observed in single-cell data, which is critical for estimating power to detect frequency shifts in cell states.

Here, we present single-cell Power Simulation Tool (scPOST), a novel simulation framework that enables the fast simulation of realistic datasets with >100,000 cells spanning hundreds of samples. scPOST models inter-sample variation in cell state frequencies and generates datasets based on (1) complex batch and sample parameters learned from real input prototype data (public or pilot data), and (2) user-specified study design factors. scPOST can model large studies based on minimal input data and allows users control over the degree of influence that covariates have on gene expression, ultimately informing optimal study design. We apply scPOST to three diverse single-cell datasets with the overall goal of guiding the design of larger studies that maximize power.

## Results

### Summary of statistical approach

scPOST comprises three steps: (1) parameter estimation from a prototype dataset, (2) simulation of datasets based on estimated parameters, and (3) power calculations from association testing on the simulated data (**Fig. 1**).

**Figure 1.**
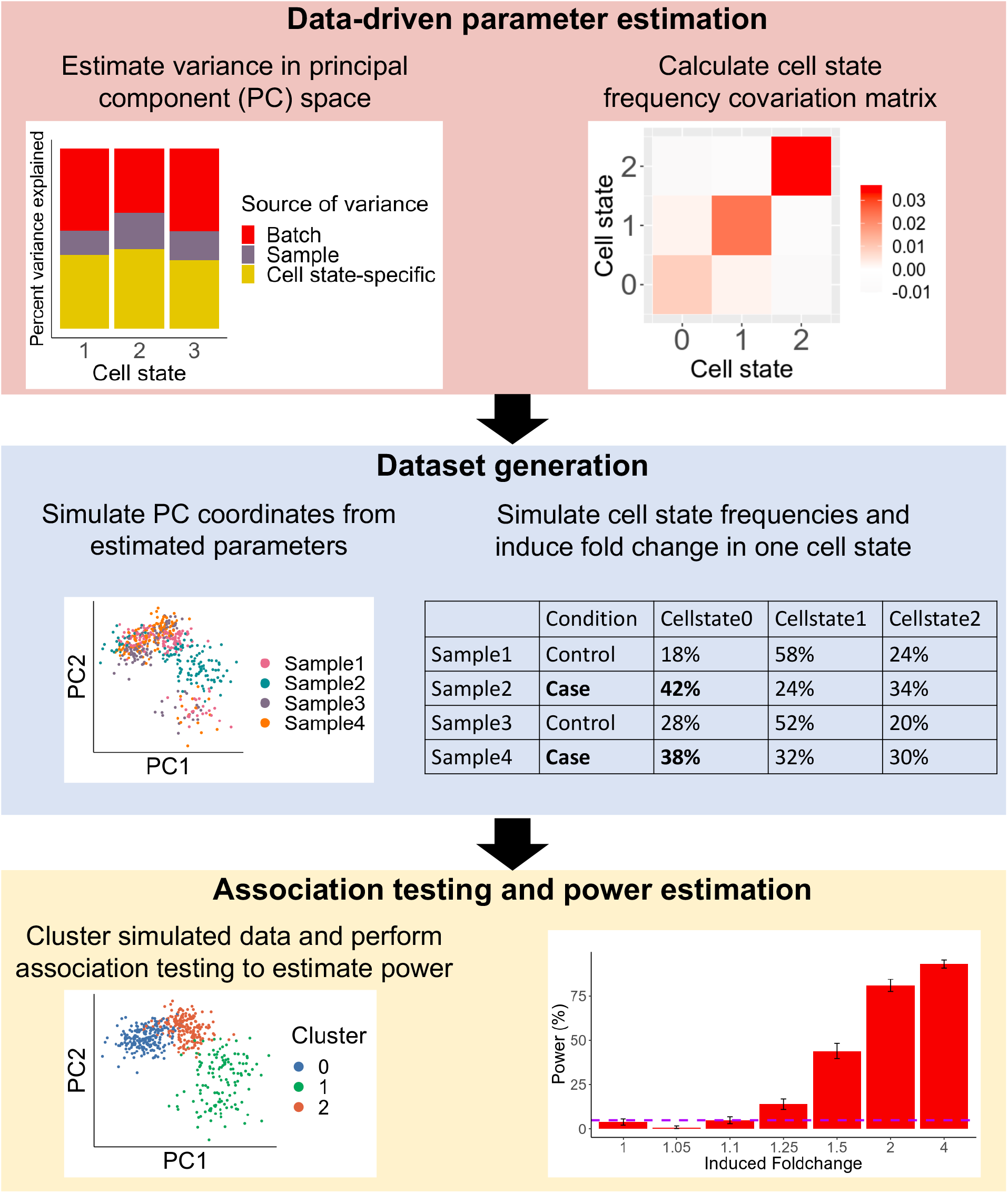
scPOST simulates single-cell datasets to estimate power to detect shifts in a cell state’s frequency across conditions. scPOST comprises 3 steps: data-driven parameter estimation, dataset simulation, and association testing. scPOST models gene expression variation (left) and cell state frequency variation (right).

scPOST relies on user-provided prototype data to estimate key parameters used for simulation. This prototype data should reflect the planned experimental setting, including the single-cell platform being used, assayed cell states, and sample type. We recommend prototype data include a minimum of 6 samples with at least 500 cells per sample so that it properly captures the inter-sample cell state and gene expression covariation of the expected data. The user can control key study design factors including: number of cell states, number of samples and cells per sample, multiplexing structure, and magnitude of simulated noise. scPOST allows users to input their own pre-determined cell states; in the following examples, we use cell states derived internally from principal component analysis (PCA)^25^.

We model gene expression in low-dimensional principal components (PCs) rather than high-dimensional measurements of individual genes to preserve gene covariation structure. For each cell state cluster, we estimate the variance in PC space captured by batch ∑_B_ and sample identity ∑_S_ by using principal variance component analysis. We use linear mixed models with batch and sample identity fitted as independent random effects to estimate ∑_B_ and ∑_S_ (**Fig. 2, Online Methods**). Next, we retrieve the cell state’s unconditioned centroid in PC space **μ**, representing the mean value along each PC in that cell state. We also obtain the residual variance ∑_C_ (the cell state’s variance in PC space not attributed to batch and sample identity), which we interpret as intrinsic variance specific to the cell state (**Fig. 2**).

**Figure 2.**
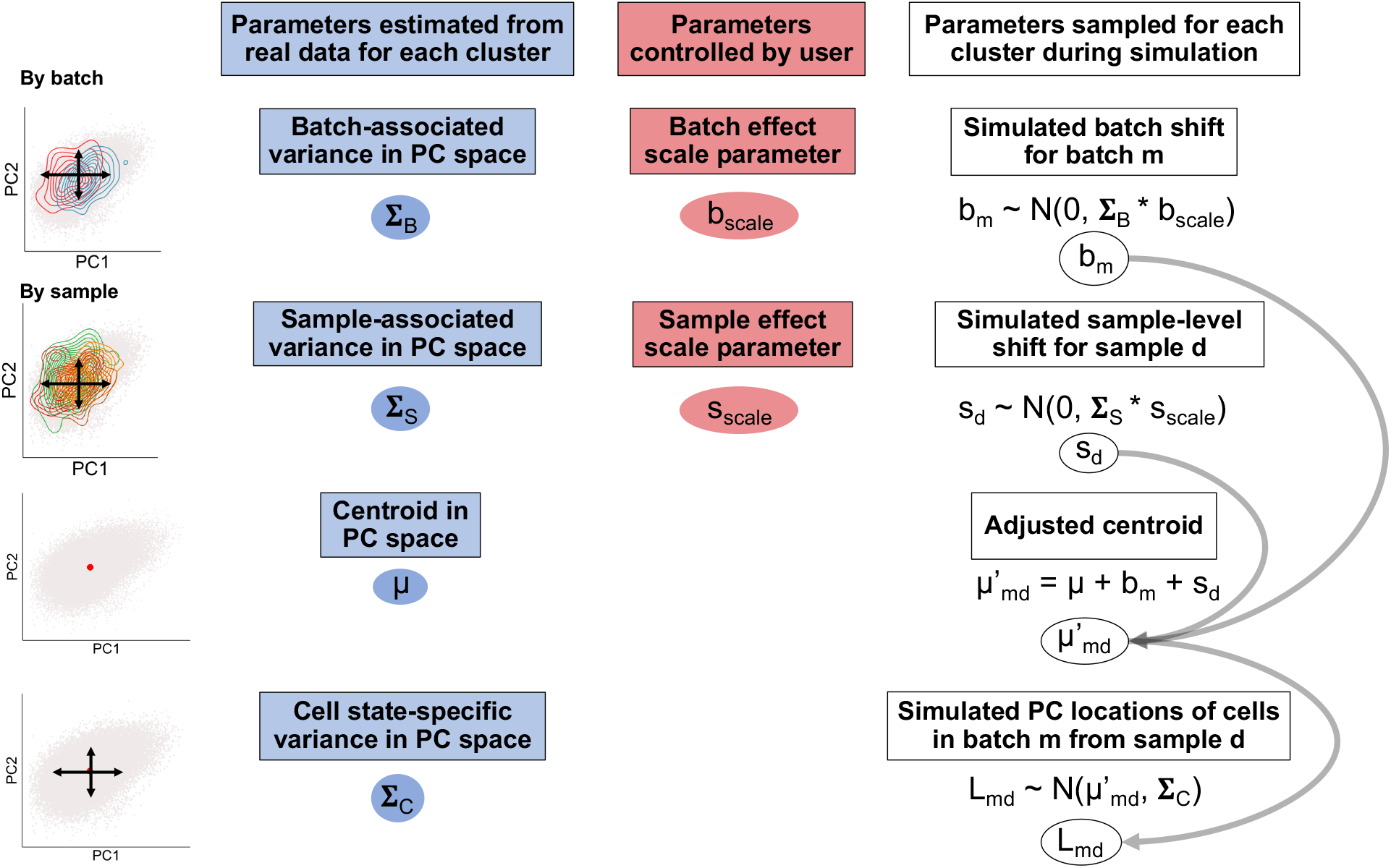
scPOST simulates gene expression by sampling principal component (PC) coordinates. scPOST generates PC coordinates for each simulated single cell. With cell state-specific parameters estimated from real data (blue) and user-controlled scaling factors (red), we simulate batch- and sample-specific shifts for each cell state cluster. These shifts are summed with the cell state specific centroid obtain an adjusted centroid; these adjusted centroids are then used to sample PC coordinates by using the residual cell state-specific variation. Thus, each simulated cell’s PC coordinate is dependent on its assigned cell state, batch, and sample.

For each cell state, we simulate random linear shifts for each batch *m* (*b_m_*) and each sample *d* (*s*_*d*_) from ∑_B_ and (∑_S_ respectively; these shifts are multiplied by user-controlled scaling factors (b_scale_ and s_scale_, respectively) to modulate the magnitude of these effects. These shifts produce an adjusted centroid **μ**’_md_ for each batch-sample pair. Finally, each cell is simulated in PC space by sampling from a multivariate normal distribution centered at **μ**’_md_ with cell state-specific covariance ∑_C_ (**Fig. 2**). Without adding large amounts of noise, our simulated dataset’s gene expression profiles closely matched the originating prototype data (**Fig. 3a-c**). Adding more noise can result in cell states mixing with each other if, for example, ∑_B_ increases (**Fig. S1**).

**Figure 3.**
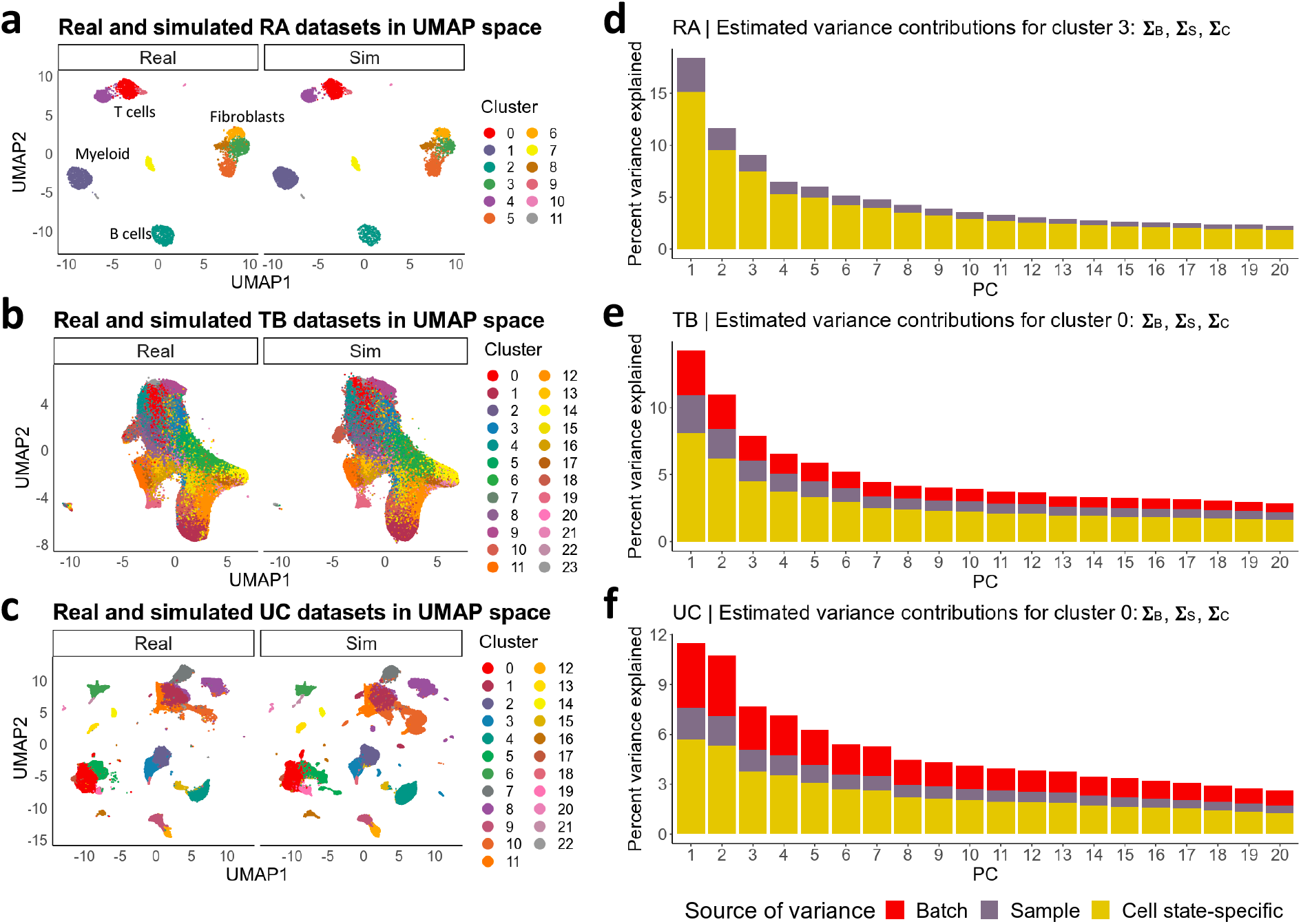
Estimated parameters from the input prototype RA, TB, and UC datasets. **a-c,** UMAP visualizations of the RA, TB, and UC datasets colored by cell state clusters. Each panel exhibits side-by-side comparisons of the real dataset and a simulated realistic (as derived from the real data) dataset which are both embedded in the same UMAP space. The simulated datasets had a similar number of cells compared to their respective original input dataset. **d-f**, Bar plots of the percentage of variance explained in each principal component (PC) in a representative cluster from each of the RA, TB, and UC datasets. “Percentage variance explained” refers to the estimated total variance of the real dataset accounted for by each PC; each bar is colored by the source of noise contributing to that variance. Cluster 3 in the RA dataset (green-colored fibroblast) is focused on in Fig. 4a and Fig. S4.

From the prototype dataset’s cell state frequency distributions, we estimate each cell state’s mean frequency **μ**_cf_ across samples and covariance with other cell states ∑_cf_ (**Fig. S2**). We simulate each sample’s cell state frequency distribution by sampling from a multivariate normal distribution parameterized by **μ**_cf_ and ∑_cf_ (**Online Methods**). To simulate shifts in cell state frequencies, the user may choose a cell state to induce a fold-change (effect size) difference between conditions. We encourage users to test a range of fold-changes, which can be based off of previous data.

To estimate power, we cluster our simulated dataset and apply Mixed-effects Association testing for Single Cells (MASC)^26^. In brief, MASC fits logistic mixed effects models on single-cell data and tests whether a full model containing case-control status explains cells’ cluster membership significantly better than a null model without case-control status. We considered successful detection when, after inducing a fold-change difference, at least one cluster had a significant differential frequency between conditions (**Online Methods**). Unless otherwise stated, we performed 500 simulations for each parameter combination and defined power as the percentage of successful simulations, e.g. 50% power indicates successful detection in 250/500 simulations.

### scPOST estimates parameters from three input prototype datasets

To demonstrate the utility of scPOST, we applied our framework to three diverse scRNA-seq datasets derived from synovial tissue (rheumatoid arthritis: RA)^10^, peripheral blood memory T cells (Tuberculosis: TB)^14^, or intestine (ulcerative colitis: UC)^15^ (**Table S1**). The RA dataset is smaller than the TB and UC datasets, both in number of cells and number of samples assayed: 5,265 cells (21 samples) versus 496,517 cells (259 samples) and 235,229 cells (30 samples) respectively. The RA and UC datasets contain many broad immune and stromal cell types while TB contains only memory T cells.

We applied a standard PCA dimensionality reduction pipeline to each dataset (**Online Methods**). For RA, we corrected the PCs for batch- and sample-level effects with the Harmony algorithm^27^ and clustered with the Louvain method^28,29^, yielding 12 transcriptionally distinct putative cell states (**Fig. 3a**). We did not correct batch effects in the TB and UC datasets to model the performance of scPOST in datasets with different levels of batch effects. We found 24 and 23 putative cell states for TB and UC respectively (**Fig. 3b-c**). Each dataset’s PC coordinates and cell states were the scPOST input for their respective power analyses.

scPOST highlights the diversity of these single-cell datasets. For example, the RA and UC datasets featured higher levels of residual cell state-specific variance, likely because they contained multiple distinct broad cell types (**Fig. 3d-f, Fig. S3**). Since the RA dataset was batch-corrected, its cell states had lower levels of batch-associated variance (relative to the intrinsic variance) compared to the TB and UC datasets; cell states with low gene expression variance should be easier to classify and associate with clinical phenotypes because they are less likely to mix with other cell states. Cell states in the RA and UC datasets exhibited higher frequency variation compared to TB (**Fig. S2**), likely because they are derived from more heterogeneous dissociated tissue compared to the less destructive collection of blood.

### scPOST estimates power benefits from expanding RA study

We first used scPOST to determine how to best expand the RA study to more reliably detect differential frequencies. In cytometry data, the original authors (along with others) identified an *HLA*^hi^ fibroblast subset that was expanded in inflamed RA samples compared to osteoarthritis (OA) controls, but could not detect this expansion in the scRNA-seq dataset due to small sample size (n = 21)^10^. We simulated realistic (derived from the RA fibroblast data) fibroblast datasets and induced a fold-change of 5 in the cluster with the highest expression of *HLA* in fibroblasts (**Fig. S4, Online Methods**). For 20, 40, or 80 samples, we simulated unbalanced study designs in the original study’s proportion with 17 case/3 control, 34 case/6 control, or 68 case/12 control samples respectively. Power was only 12% for 20 samples but increased to 29% for 40 and 75% for 80 (**Fig. 4a**). We repeated these power analyses, but this time with study designs containing an equal number of cases and controls (**Online Methods**). In this context, power dramatically increased to 60% at 20 samples, 89% at 40, and a full 100% at 80 (**Fig. 4a**). Thus, we determined the number of samples needed to maximize power and estimated the benefit of a balanced study design.

**Figure 4.**
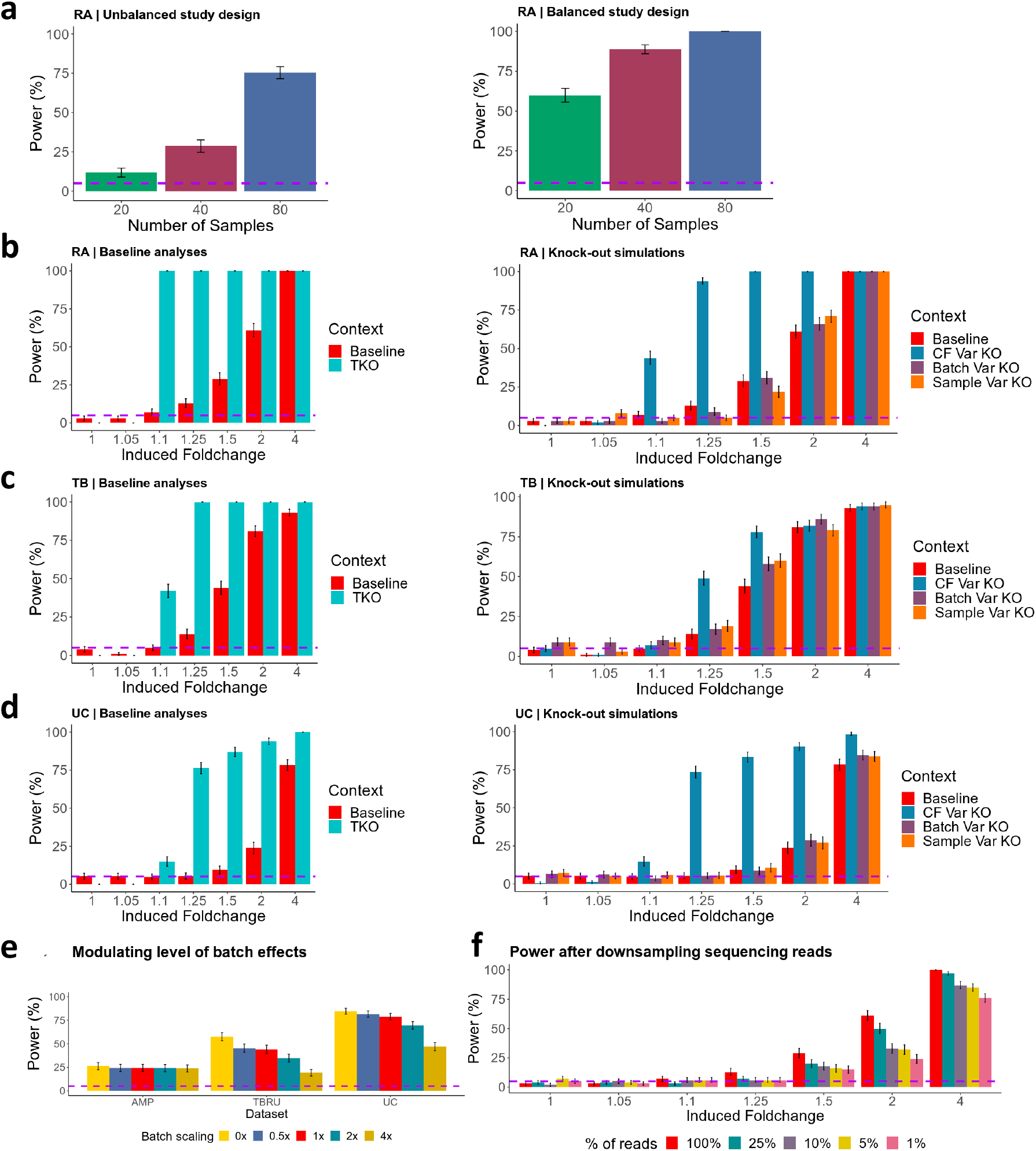
scPOST estimates power from multiple study designs. Each bar represents power estimation from (n = 500) simulations. In all panels, error bars represent 95% binomial proportion confidence intervals and the dotted horizontal purple line represents 5% power. **a,** Results from increasing the size of the RA study. **b-d,** Baseline and “knock-out” analyses for the RA, TB, and UC datasets. The baseline context features realistic levels of cell state frequency variation and gene expression variation, while knock-outs remove the respective variation (TKO = triple-knock-out where all three sources were removed, CF = cell state frequency). **e,** Batch scaling simulations: 0x indicates no batch variation, while 4 indicates 4x times the level of estimated realistic (1x analogous to baseline in **b-d**) batch variation. Induced fold-change of 1.5 in RA and TB settings and fold-change of 4 in UC. **f,** Power calculations from datasets derived from downsampling the AMP read data (% of reads refers to percentage of downsampled reads compared to the original data, 100% analogous to baseline).

### Baseline power analyses estimate realistic power

We then performed a “baseline” power analysis for each dataset to illustrate power under realistic conditions; we used these results to benchmark study design changes. We simulated single-cell datasets with realistic (derived from input data) cell state frequency variation and batch- and sample-level gene expression variation (**Online methods**). Each simulation contained 50 case and 50 control samples, 500 cells per sample, and 4 samples per batch. In each simulation, we induced a fold-change difference in one uniformly random cell state so that our power estimates are an aggregate of all cell states.

In each experimental setting (RA, TB, UC), we simulated a dataset and estimated power to detect differential abundance over a range of fold-changes: 1 (no change), 1.05, 1.1, 1.25, 1.5, 2, and 4, corresponding to a 0%, 5%, 10%, 25%, 50%, 100%, and 300% respective increase in a cell state’s frequency in cases versus controls. Power in the TB setting was generally greater or equal to power in the RA and UC settings (**Fig. 4b-d left**). This difference reflects more modest estimated cell state frequency variances in the TB data compared to RA and UC.

### Estimating upper limits on power

Next, we wanted to estimate how intrinsic gene expression variation leading to imprecise clustering might limit power in the absence of other sources of noise. Since the TB and UC settings each have many transcriptionally similar fine-grained states within broad cell types, we predicted lower power estimates compared to RA, whose cell states are more transcriptionally discrete.

Accordingly, we performed analyses in which we simultaneously “knocked out” three sources of noise (triple knock-out = TKO). We assumed no cell state frequency variation and no batch-associated or sample-associated gene expression variation so that all noise is driven by intrinsic gene expression variation (**Online Methods**). If there were absolutely no variation, we would have 100% power to detect a frequency shift of any magnitude; therefore, any reduction of power observed in the TKO context is due to intrinsic gene expression variation. In all three experimental conditions, we had 0% power to detect an extremely subtle fold-change of 1.05, indicating that intrinsic gene expression variation limits power even in the absence of other sources of noise (**Fig. 4b-d**). However, we estimated 100% power at fold-changes as little as 1.1 and 1.25 in the RA and TB settings respectively.

### Identifying which source of variation most affects power

We next wanted to determine which of the three TKO sources of variation had the most effect on power. Accordingly, we performed simulations in which we only “knocked out” individual sources of variation while leaving the other sources of variation at levels derived from the prototype dataset (**Online Methods**).

First, we removed cell state frequency variation so that each simulated sample had the same cell state frequency distribution before inducing a fold-change in a cluster. This resulted in increased power, especially in the RA and UC settings (**Fig. 4b-d right**). Considering the RA and UC cell states tended to have higher variation in frequencies compared to those in the TB dataset, it is unsurprising that the increases in power were higher in these settings.

Next, we removed only batch-associated variation in gene expression (**Fig. 4b-d right**). We observed minimal increases in power at most fold-changes for all three settings. Finally, we removed only sample-associated variation in gene expression and again observed modest changes in power for each fold-change in each setting (**Fig. 4b-d right**). These results suggest that cell state frequency variation has the most effect on power.

### Modulating the magnitude of batch-associated transcriptional variation

Batch effects can be reduced by investing time and energy into improving protocols and reproducibility; however, the value of this investment is not always clear. To quantify how much decreasing batch effects might increase power, we performed analyses where we modulated the levels of batch-associated variation (**Online Methods**). We used a scaling factor to scale prototype-derived estimates of batch-associated variance by 0x, 0.5x, 1x, 2x, or 4x. The batch-corrected RA setting was largely resistant to high multipliers of estimated batch-associated variance while the TB and UC settings showed decreased power at higher multipliers of batch-associated variance (**Fig. 4e**).

### Extremely low sequencing depth decreases power

Sequencing depth is an important property of single-cell studies because it not only affects the detection of transcripts, but also the precision with which cells are classified and thus, power. To measure how reduced sequencing depth affects power, we first downsampled the RA dataset’s sequencing reads (**Online Methods**). Compared to the 7,300 mean reads of the original dataset (**Fig. S5**), we created downsampled datasets with 1,825, 730, 365, and 73 mean reads (25%, 10%, 5%, and 1% of the original sequencing depth, respectively). For each downsampled dataset, we applied a standard PCA pipeline. UMAP visualization suggests that reducing read depth to 10% or lower results in cell state mixing (**Fig. S6**). The PCs obtained for each downsampled dataset and the clusters obtained from the original dataset were input into scPOST.

We observed that at most fold-changes, there were minimal differences in power between the original dataset and the dataset with 25% of reads (**Fig 4f red/teal**). However, when downsampling to 10% of reads or lower, we noticed power loss at fold-changes above 1.5. These results suggest that reducing sequencing depth to extremely low values can eventually confound cell state classification and power.

### Multiplexed study designs can decrease batch effects and increase power

Multiplexing samples by running multiple samples in a single batch can reduce the number of batches in a study, and thus the impact of batch effects. We sought to estimate how power is affected by reducing the number of batches (by increasing the number of samples per batch). In the TB setting, we modulated the magnitude of batch-associated variance in addition to reducing the number of batches (**Online Methods**). At realistic (1x) levels of batch variance, reducing the number of batches resulted in minimal power changes. In contrast, when we assessed power at 4x the realistic batch-associated variance, we observed that multiplexing to reduce the number of batches rescues power to levels observed at 1x (**Fig. S7**). While effective, this strategy requires a small number of batches with large numbers of samples and cells, which may not be feasible in all experimental settings.

We then investigated another multiplexing scheme that retained the same number of cells per batch and total samples, but distributed samples across multiple batches. As a control, we assessed a sequential study design with realistic parameters consisting of 100 samples (2000 cells each) placed into individual batches. This was compared with a multiplexing scheme that still contained 100 samples of 2000 cells each across 100 batches, but each batch contained 500 cells from 4 different samples (**Fig. S8, Online Methods**). Multiplexing yielded noticeable improvements, especially at higher multipliers of batch effects (**Fig. 5a**). These benefits are likely due to the multiplexing design enabling better estimation of batch effects on individual samples.

**Figure 5.**
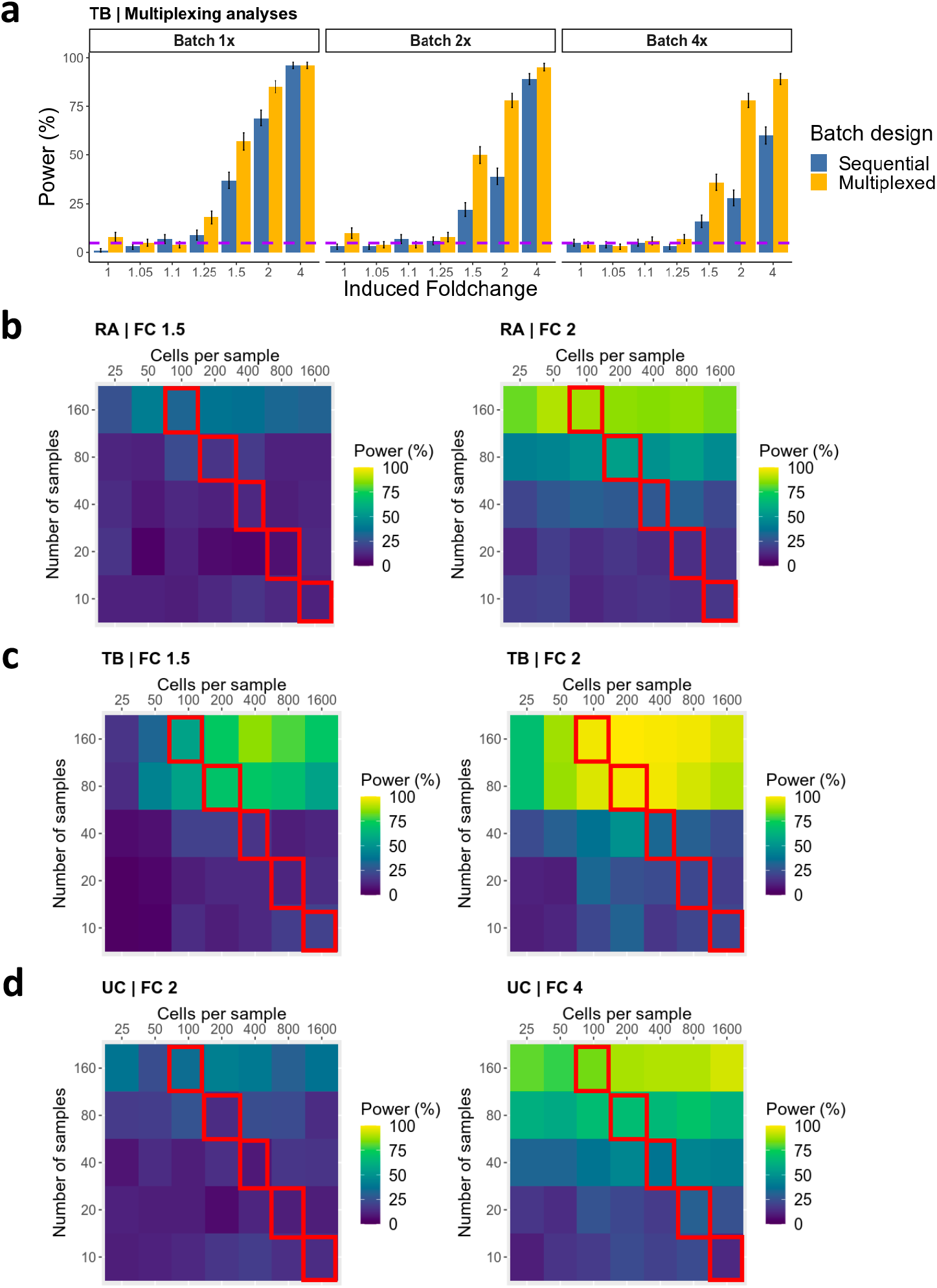
Measuring benefits of multiplexing and increasing dataset size. **A,** Power estimates comparing a sequential versus a multiplexed study design (Fig. S8). Simulations with these study designs were also run with higher scaled levels of batch effects (2x and 4x) compared to the realistic level of batch effects (1x = baseline). Each bar represents power estimation from (n = 500) simulations. Error bars represent 95% binomial proportion confidence intervals and the dotted horizontal purple line represents 5% power. **B-D**, Power calculations across different ranges of dataset sizes at fold-changes of 1.5, 2, or 4. Each tile represents power estimation from (n = 100) simulations. The total size of a dataset is determined by the number of samples and cells per sample. Simulations were performed in a realistic context (derived from dataset-driven parameter estimates) with varying numbers of samples/cells per sample. Tiles across a top left-bottom right diagonal have an equivalent number of cells, and tiles outlined in red all have 16,000 total cells.

### Increasing the number of samples improves power more than increasing cells per sample

Collecting a large number of samples can be difficult, especially in human studies, due to availability and cost. Likewise, increasing the number of cells per sample can be prohibitive because of protocol limitations and constraints on the size of each sample, leading to important tradeoffs in cost and power. Thus, we investigated whether more power is gained by increasing the number of total samples versus increasing the number of cells per sample.

In each of the three experimental settings, we simulated datasets with the same parameters as our baseline analyses (**Fig. 4b-d**) while varying the number of samples and cells per sample. We simulated datasets across two ranges: 10, 20, 40, 80, or 160 samples and 100, 200, 400, 800, and 1600 cells per sample. For simulations with 16,000 total cells (**Figure 5b-d red outline**), we simulated 10, 20, 40, 80, and 160 samples with 1600, 800, 400, 200, and 100 cells each, respectively. In all three experimental settings, increasing the number of samples yielded noticeably higher increases in power compared to increasing the number of cells per sample for the same total number of cells. We observed a similar pattern across all tested fold-changes (**Fig. S8-10**).

## Discussion

Here we present scPOST and demonstrate its utility by simulating thousands of large single-cell datasets to investigate how different study design choices affect power to detect shifts in cell state frequencies across conditions. Power estimates for single-cell studies of clinical samples are essential to inform optimal allocation of limited resources, e.g. patient samples and single-cell assays. scPOST is particularly useful because it estimates power from a user-defined prototype dataset, which may be a public or pilot dataset (such as from the Human Cell Atlas^30^) that reflects the planned experimental setting.

In a powerful use case, we used scPOST to determine how to expand the 21-sample RA study to more robustly detect an expansion of *HLA^hi^* fibroblasts. We found we could significantly increase power by adopting a balanced study design and obtaining 20 more samples. Given that we (and others^24,31^) found that shallower sequencing or assaying fewer cells per sample does not generally compromise power, it is more effective for investigators to allocate these resources to obtaining more samples.

Across three diverse experimental conditions, we consistently found that three factors substantially influenced overall power: the number of independent samples, a cell state’s frequency variation across samples, and the magnitude of batch effects. In contrast, we found that acquiring more cells per sample and deeper sequencing had more modest effects on power.

We note that minimizing batch effects has benefits besides power, since batch effects can result in the identification of spurious cell states. We observed that a multiplexing scheme that splits samples across multiple batches can provide significant benefits to power by reducing batch effects. If it is more difficult for a study to obtain more samples, but the samples are big enough to be split into multiple batches, utilizing this multiplexing scheme may be a way to increase the study’s power.

We choose to simulate gene expression as PC coordinates; this precludes these datasets from being analyzed by tools that require raw gene expression data. Nevertheless, scPOST allows users to output the simulated datasets, which can be analyzed with tools that utilize PC coordinates or graphs derived from PC coordinates, such as clustering algorithms or trajectory analysis^32,33^.

We recognize that association testing can be conducted on various groupings of cell states obtained from different clustering resolutions. Low-resolution groupings may mask signal in smaller cell states, while high-resolution groupings may break up signal in larger cell states. We recommend users try a range of clustering resolutions.

scPOST currently operates using a low-dimensional PCA embedding of cells. With multimodal technologies such as CITE-seq^5^ becoming more available, analyses of these new data types may include dimensionality reduction with alternative methods, such as canonical correlation analysis (CCA)^10,14,34^ and nonlinear embeddings^35,36^. Simulating new types of data in the context of these alternative tools, such as simulating canonical variate coordinates instead of PC coordinates, represents a possible extension of scPOST. Another potential extension is associating cell state frequencies with alternative variables beyond case-control status, including continuous variables (*e.g.* polygenic risk score).

We envision that scPOST will be applied to representative public datasets or pilot studies in order to predict power of future studies. Alternatively, our tool can be used to determine the range of effect sizes a completed study is actually powered to detect. As single-cell studies become larger and shift towards comparing conditions, we expect scPOST to be broadly useful for investigators in many experimental settings for defining optimal study designs.

## Online Methods: single-cell Power Simulation Tool (scPOST)

### Overview

The goal of scPOST is to provide a flexible tool to simulate single-cell datasets that approximate an experimental context. scPOST uses input prototype datasets (e.g. a public dataset or pilot dataset) to estimate characteristic qualities of an experimental setting, such as the variation in cell state frequencies across samples and the gene expression covariation structure of cells likely to be measured in the setting. By modifying the study design of the generated datasets, investigators may use scPOST to predict how specific study design choices might affect the results of a single-cell experiment. We envision scPOST will be helpful for investigators in defining the design of their studies so that they may maximize their power to detect shifts in cell state frequencies between conditions (e.g. expansion of a cell state in case samples or stimulated samples).

Here, we present scPOST which comprises three steps: (1) parameter estimation that takes a prototype dataset as input, (2) simulation of new single-cell datasets based on the estimated parameters, and (3) power calculations from performing association testing. The user is able to customize the construction of the simulated dataset in several ways including: the number of total cells, the number of total samples, the number of cells per sample, the number of batches, multiplexing structure, and the magnitude of noise that contributes to the variation of cell state frequencies or cell state gene expression. In this paper, the association testing we pair with scPOST is a test for differential abundance: associating a cell state with shifts in frequencies between conditions (e.g. case-control status). The simplest use case we present in the paper is testing our power to detect an expansion of a cell state cluster in case samples, but scPOST allows investigators to modulate many aspects of the study design to explore different scenarios.

We allow for two main types of output: simulated datasets and power calculations from testing for differential abundance. The results presented in this paper are power calculations from testing for differential abundance, but if users want to perform different association tests on the simulated datasets, they may retrieve the datasets instead. Implementations of scPOST are available as part of an R package at https://github.com/immunogenomics/scpost, along with several of the input prototype datasets showcased in this paper. The following sections explain the procedures used in the scPOST framework. We begin with how we test for differential abundance and estimate power, then proceed into how we simulate datasets.

## Step 3: Power estimation from association testing

### 1.1 Testing cell state clusters for differential abundance with MASC

The output of **Step 2** is the principal component (PC) coordinates of a generated single-cell dataset, which are a representation of the gene expression space. Here, we generate new cluster labels for our simulated cells by creating a shared-nearest neighbor (SNN) graph (k = 30) and using Louvain’s method of community detection at the same resolution used in the original input prototype dataset (resolution controlled by user). From this procedure, we obtain cell state cluster assignments for each of our simulated cells, which we then use to test for associations.

We test whether cell state clusters have differential abundance between two conditions, which we refer to as case vs. control. In conventional RNA sequencing (RNA-seq), it is common to apply Welch’s unequal variance t-test to test whether a cell state cluster’s mean frequency in cases is significantly different from the mean frequency in controls. It is not enough to count the number of case cells compared to controls, as it is often the case that cell state frequency variation results in many cells of interest being derived from one or two samples. However, in single-cell RNA-seq, this statistical test has shown to be underpowered. Here, we use an alternative method, Mixed-effects Association testing for Single Cells (MASC)^26^, which is a higher-powered differential abundance test that utilizes logistic mixed effects models to test whether a cell state cluster’s identity is associated with its condition status. MASC allows us to account for potentially confounding single-cell covariates by fitting null and full models such covariates. For all of our results, the null and full models used as input for MASC are as follows:

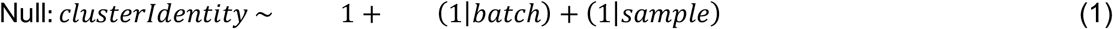

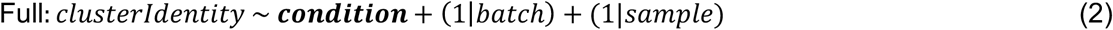

Where *clusterIdentity* refers to the cell state cluster assignments obtained from clustering the simulated dataset, *condition* refers to the category the simulated cell belongs to (e.g. case or control), *batch* refers to the batch the cell belongs to, and *sample* refers to the sample the cell belongs to.

### 1.2 Power estimation

For each simulation, we test all cell state clusters for differential abundance. Here, we considered a simulation to have successful detection if at least one cluster had a significant differential abundance between conditions. Significance is defined by the Bonferroni-corrected threshold of p < 0.05/*k*, where *k* is the number cell state clusters we test. We note that it is often the case that the cell state we induce a fold-change into contributes to one of these clusters, but we do not require it to, because changing the frequency of a cell state in a sample inherently changes the frequency of other cell states in the sample (due to the number of cells in that sample being fixed). Furthermore, cell states can covary with others, which can lead to a cell state exhibiting frequency shifts between conditions, even though we did not induce a fold-change into that cell state. We performed many simulations (either 100 or 500) for a combination of parameters and defined power as the number of simulations in which we successfully detected differential abundance (e.g. 50% power means we successfully detected differential abundance in half of the simulations). The error bars in the power plots are 95% binomial proportion confidence intervals.

## Step 2: Generating a single-cell dataset

In our simulations, we use an input prototype dataset *X* to generate a new single-cell dataset *S* in the form of a cell by PC coordinate matrix. As part of dataset generation, we also assign single cells to a sample and batch. We focus on two general aspects of variation in a single-cell dataset: (1) the variation in gene expression, which we represent with PC coordinates and (2) the variation in cell state frequencies across samples.

### 2.1 Representing a single-cell in PC coordinates

Principal components derived from principal component analysis (PCA) on a cell by gene matrix, can be interpreted as summaries of gene expression and represent data structure in the context of covariation between genes. PCA is commonly used in dimensionality reduction pipelines for single-cell analyses. The PC embeddings (values) for each cell represent its location in PC space and gene expression patterns relative to other cells. Cells closer to each other in PC space generally have more similar gene expression profiles than cells further away from each other, which means the PC coordinates of a single-cell dataset also encode covariation structure between genes. Due to this, we represent the gene expression of a cell by their PC coordinate; we estimate variation in PC space in parameter estimation (**Step 1**), and we generate PC coordinates for each simulated cell (**Step 2**). The generated PC locations for our simulated cells *L* are used for clustering and power estimation (**Step 3**).

### 2.2 Simulating PC coordinates for a cell state

Our simulation approach assumes that each cell state has different levels of variation. Furthermore, we assume that the gene expression of each cell state is affected by batch or sample effects (derived from the batch the cell was run in or the sample origin respectively) differently.

For simulated cells, we assign initial cell state identities that are analogous to the cell states in the original input dataset *X*; these assignments are derived from simulated cell state frequency distributions that we define in **section 2.4**. Based on the simulated cell’s assigned cell state, we generate PC coordinates. If *c* is the number of cells in a simulated dataset and *e* is the number of PC dimensions we simulate, we simulate a matrix of PC coordinates whose dimensions are *c* × *e*.

For each cell state cluster, we assume that its PC coordinates are distributed around some *e*-dimensional centroid **μ** with covariance *∑*_C_. In our simplest generative model without batch or sample effects, we simulate an *e*-dimensional vector of PC coordinate for each cell based on its assigned cell state. To generate *L*, we sample from a multivariate normal distribution parameterized with mean **μ** and covariance *∑*, giving *L* the following distribution:

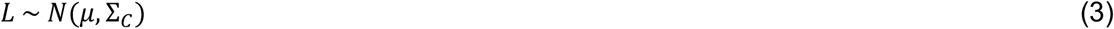

### 2.3 Simulating technical effects (batch and sample-level effects)

Variation in gene expression may influence our ability to detect differential abundance, as high variation may result in misclassification of cell states via clustering algorithms. Because PCs are a summary (linear projection) of gene expression patterns, variation in PC space represents variation in gene expression. Here, we use linear mixed models to explicitly model the magnitude of batch-associated variation in PC space and sample-associated variation in PC space. We assume that each cell state is affected by batch and sample effects in its own way, so we model and simulate a batch/sample effect for each cell state we simulate.

We simulate batch and sample-level effects by using parameter estimates retrieved from **Step 1**(**Section 3.1**). These effects take the form of linear shifts that are summed with a cell state’s centroid. We assume that batch linear shifts have a multivariate normal distribution with mean 0 and covariance *∑*_B_ (which represents the within-cell state variance of cells in PC space contributed from batch effects). Similarly, we assume that sample linear shifts have a multivariate normal distribution with mean 0 and covariance *∑*_S_ (which represents the within-cell state variance of cells in PC space contributed from sample effects).

In order to modulate the magnitude of these batch and sample linear shifts (and thus the magnitude of the batch/sample effects), we use scale factors, b_scale_ and s_scale_ respectively. If b_scale_ is set to 0, there will be no batch linear shift (equivalent to 0 batch-associated effects in the simulated dataset). The same is true for s_scale_. These scale factors modulate the covariance of the multivariate normal distribution we sample from. Thus, our simulated batch/sample linear shifts for batch *m* or sample *d* respectively have the distributions:

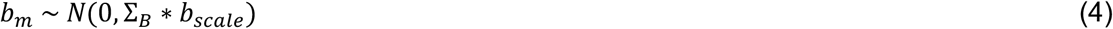

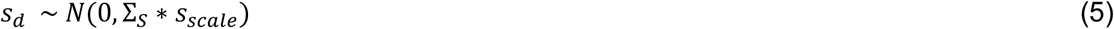

Each cell state’s batch and sample linear shifts are summed with its respective cell state centroid in order to create an adjusted centroid that is dependent on the cell’s state, batch identity, and sample identity. Thus, a cell state’s adjusted centroid for cells from batch *m* and sample *d*, **μ**’_md_, is formulated as:

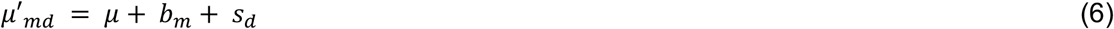

In our generative model that incorporates batch and sample-level effects on gene expression, a cell state’s adjusted centroid and its respective cell state covariance *∑*_C_. Thus, a cell state’s PC locations for cells from batch *m* and sample *d*, *L*_*m,d*_, are distributed as:

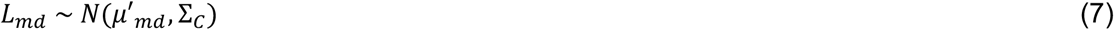

### 2.4 Simulating cell state frequency distributions

Since the power to detect differential abundance in an experiment is also dependent on the frequency distributions across the samples, it is important for simulations to facilitate the simulation of cell state frequency distributions that reflect the cell state frequency variation observed in the experimental setting (which can be relatively high, especially in human data). From input prototype data, we empirically estimate a *k*-length cell state mean frequency (CF) distribution **μ**_cf_ (whose elements contain the observed mean frequency of a cell state across all of the samples in the dataset) and its corresponding variance-covariance matrix *∑*_cf_; see **Section 3.2** for the estimation procedure.

For each sample (total number set by user), we generate a cell state frequency distribution, which is defined as the proportion of each cell state in a sample. The number of cell states generated in these distributions is dependent on the number of cell states in the original input data.

Since we work with frequency distributions and variation in those frequencies, we transform **μ**_cf_ and *∑*_cf_ into log-space (so that we do not sample negative frequencies). We then sample from a multivariate normal distribution parameterized by mean **μ**_cf_ and covariance *∑*_cf_; we allow users to scale the amount of cell state frequency covariation is generated via a scaling parameter cf_scale_. If cf_scale_ is set to 0, the simulated samples will all have the exact same cell state frequency distributions. The resulting *k*-length vector is then transformed back into linear space. The cell state frequency vector for a sample *F* is thus distributed as:

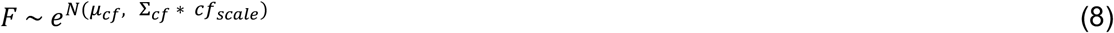

Finally, each value in *F* is divided by the sum of all values in *F* to create a probability distribution. Each sample’s generated probability distribution is used to determine the cell state assignments for cells assigned to that sample. Samples are pre-assigned to batches based on the user-defined batch structure. We provide functions that help users split samples into batches (based on the R function “split”).

### 2.5 Inducing a fold-change difference in a cell state between conditions

Here, we define a fold-change in a cell state as the ratio of cells in one condition compared to the other (e.g. a fold-change of 4 means the number of case cells is 4x compared to the number of control cells). In our simulations, we induce a user-defined fold-change difference in one cell state. This fold-change difference is what we wish to detect. To perform the fold-change difference in a cell state between conditions, we alter all of the CF distributions generated for all samples in one condition (here, we alter the case samples). We first scale the frequency of the selected cell state by the magnitude of the fold-change we wish to induce, which gives us the new frequency for the cell state of interest and remaining probability mass. The frequency for the cell state is capped at 1 (which would mean all cells from that sample come from this cell state). The frequencies for the remaining clusters are then divided by their sum (the sum of the frequencies of the remaining clusters), and then scaled by the remaining probability mass. If there is no CF variation cf_scale_ = 0, then the number of case cells will outnumber the control cells exactly by the fold-change induced.

## Step 1: Parameter estimation

To generate a new single-cell dataset *S* in the form of a cell by PC matrix, we estimate several key parameters from the original input prototype dataset. The PC embeddings (received from the PCA) cell state cluster assignments are used as input for parameter estimation (we treat clusters as cell states).

### 3.1 Using linear mixed models to estimate gene expression variation in principal component space

We focus on variation in PC space, which represents variation in gene expression. To deconvolute the different sources of this variation, we use principal variance component analysis (PVCA). While we focus on the contributions that batch- and sample-specific effects have on gene expression variation, we note that more or different covariates may be fit in these models. This is a possible extension that users can make, but they would need to augment the generation of the single-cell datasets to include the new/different covariates, as well as the model they use to test for differential abundance. Linear mixed effects models fit equations of the form:

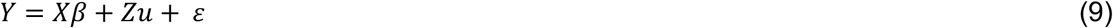

Where the unconditional distributions of random effects designated by *u* are:

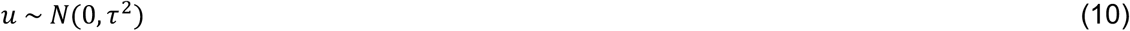

The *τ*^2^ parameter of a random effect represents its estimated variance contribution. Again, we assume that e each cell state cluster has a different level of batch-associated and sample-associated variation. Thus, for each cell state *k*, we fit the following formula for each PC (we use a default of 20 PCs) with the R function “lmer” from the “lme4” package:

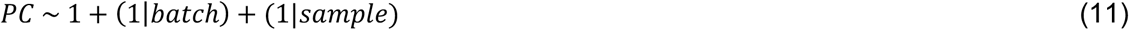

Because we fit random models for each PC for each cell state, we fit *k* × *e* models. After fitting, we extract following elements from each model:

**μ**_k_: The fixed effect intercept of a model, representing the unconditional mean value in PC space. The mean value for all PCs of a cell state represent its centroid
**∑**_B_: The estimated *τ*^2^ value for the batch random effect, representing the contribution batch effects have on gene expression variation in a cell state
**∑**_S_: The estimated *τ*^2^ value for the sample random effect, representing the contribution sample origin has on gene expression variation in a cell state
**∑**_C_: The estimated residuals, representing intrinsic cell state-specific variation in a cell state (the variation not contributed from our explicitly fitted covariates). This variation may contain variance associated with other covariates we did not fit.

For each cell state, we enforce *∑*_B_ and *∑*_S_ to be diagonal variance-covariance matrices where the diagonal elements are the elements of *∑*_B_ or *∑*_S_ respectively. The residual values *∑*_C_ are placed into an *e* × *e* variance-covariance matrix for each cell state.

### 3.2 Estimating cell state frequency variation

For each cell state, we estimate its mean frequency and variation across samples. The variation can be quite high, especially if the samples are from humans. The observed cell state frequency variation is likely a result of the differences between the true distribution of cell states from the sample’s origin and technical sampling variation induced by obtaining and processing the samples. Here, we estimate the full variation and do not distinguish between different sources of variation.

To calculate the mean and variation of cell state frequencies, we first count the frequency of each cell state in each sample, and then increment each frequency by a pseudo-count of 1. Within each sample, the frequencies are transformed into proportions by dividing each frequency by the total number of cells within that sample. The mean of these proportions across all samples is represented by **μ**_cf_.

During dataset generation (**Step 2**), we sample CF distributions in log-space. Thus, **μ**_cf_ is transformed into log space. Furthermore, the aforementioned proportions are then log-transformed, which we then use to calculate a variance-covariance matrix of the cell state frequencies *∑*_cf_ in log space.

## Analysis Details

As outlined in **Table S1**, the rheumatoid arthritis (RA), tuberculosis (TB), and ulcerative colitis (UC) datasets were generated in three separate studies with diverse settings^10,14,15^. The scRNA-seq RA data was derived from synovial tissue obtained from joint replacement procedures or ultrasound-guided biopsies. The scRNA-seq TB data was derived from peripheral blood mononuclear cells (PBMCs) in blood. The scRNA-seq UC data was derived from intestinal tissue obtained from biopsies.

### 4.1 Pre-processing scRNA-seq data

We applied standard scRNA-seq pre-processing steps for the RA, TB, and UC datasets^25^ independently. We removed cells whose total reads consisted of >20% mitochondrial reads. For the TB dataset, we used only memory T cells; we removed gamma delta T cells. For the UC dataset, we used only cells from healthy colitis samples taken from healthy donors and inflamed colitis samples taken from case donors; we removed cells from uninflamed colitis samples taken from case donors. The final number of cells used for our analyses is 5,265, 496,517, and 23,229 cells for the RA, TB, and UC datasets.

We then applied standard log(CP10K) normalization for gene counts using a scale factor of 10,000; for the RA dataset, we used log2 to mimic the original author’s analysis, while we used natural log for the TB and UC datasets. We found the top 2,000 variable genes using the “vst” option of the Seurat function FindVariableFeatures. Finally, we z-scored the genes.

### 4.2 Principal components analysis and batch correction

We performed PCA with the R function “rsvd” from the “rsvd” package. As is standard, we weighted the resulting eigenvectors by the calculated eigenvalues to retrieve the PC embeddings. For reproducibility, we provide the PC embedding matrices, and the meta tables that we used for the RA and UC datasets in the form of objects included in the scPOST package (ra_HarmObj and uc_Obj respectively). We also provide the gene expression for QCed cells in separate objects (geneExpr_ra and geneExpr_uc respectively).

We performed batch correction on only the RA dataset with the Harmony batch correction algorithm. We used the HarmonyMatrix function from the “harmony” package with default parameters except for the following: vars_use = c(“plate”, “sample”), do_pca = FALSE, and npcs = 20.

### 4.3 Clustering

We performed clustering by building a shared nearest-neighbor (SNN) graph (k = 30) from the PC embeddings, and then using Louvain’s method for community detection. We used resolutions of 1.2, 2.0, and 0.4 for the RA, TB, and UC datasets respectively. We used the same respective resolutions for datasets generated from the RA, TB, and UC datasets.

### 4.4 Fitting linear models

To fit our linear mixed models, we used the “lmer” function from the “lme4” package with the “nloptwrap” optimizer.

### 4.5 *HLA^hi^* fibroblast power analysis

In these analyses, we focused only on the 1,884 fibroblasts contained in the RA dataset. With only these fibroblasts, we performed the same PCA and clustering steps we applied to the other datasets. We provide the PCA embedding matrix, and meta table for these fibroblasts in the (ra_FibObj).

For each generated dataset, we assigned samples (250 cells each) into batches equally so that each batch contained 4 samples each (accomplished with our provided “distribSamples” function). We induced a fold-change of 5 in the *HLA^hi^* fibroblast cluster (which is cluster 0, index 1 in the fibroblast-only data; cluster 3, index 4 in the whole RA data), and set b_scale_, s_scale_, and cf_scale_ all equal to 1. For the unbalanced study designs, we set the following sample splits: 17case/3control for 20 samples, 34case/6control for 40 samples, and 68case/12control for 80 samples. For balanced study designs, we set the following sample splits: 10case/10control for 20 samples, 20case/20control for 40 samples, and 40case/40control for 80 samples. We ran 500 simulations for each combination of parameters.

### 4.6 Baseline power analyses

For the baseline power analyses (realistic), we assigned 100 (50 case, 50 control) samples (500 cells each) into 25 batches equally with the “distribSamples” function so that each batch contained 4 samples each. We induced the following fold-changes in case samples: 1 (no fold-change), 1.05, 1.1, 1.25, 1.5, 2, and 4. For each simulation, we induced fold-changes into a randomly chosen cluster. We set b_scale_, s_scale_, and cf_scale_ all equal to 1. We ran 500 simulations for each combination of parameters.

### 4.7 Triple-knockout (TKO) power analyses

For the TKO analyses, we used the exact same study design as the baseline analyses, but we set b_scale_, s_scale_, and cf_scale_ all equal to 0. We ran 500 simulations for each combination of parameters.

### 4.8 Singular knock-out power analyses

For singular knock-out analyses, we used identical study design parameters as the baseline analyses. However, for the cell state frequency variation knock-out (CF Var KO), we set cf_scale_ to 0 (but kept b_scale_ and s_scale_ equal to 1). For the batch variation knock-out (Batch Var KO), we set only b_scale_ to 0. For the sample variation knock-out (Sample Var KO), we set only s_scale_ to 0. We ran 500 simulations for each combination of parameters.

### 4.9 Modulating levels of batch effects

For these analyses, we used the same study design as the baseline analyses. However, we tested a range of b_scale_ values: 0, 0.5, 1, 2, and 4. The results we reported are for induced fold-changes of 1.5 in the RA and TB settings and an induced fold-change of 4 in the UC setting. We ran 500 simulations for each combination of parameters.

### 4.10 Downsampling the RA dataset and subsequent power analyses

To downsample the RA dataset, we took the raw sequencing read data from the RA dataset and performed binomial random binomial draws for each gene count using the “rbinom” function. The binomial distribution in this model is parameterized by *n* and *p*, where *n* is equal to the original gene count, and *p* is equal to the percentage of reads we want to sample (e.g. 25%). In R, this would be rbinom(1, size = *n*, prob = *p*). We downsampled datasets to have 25%, 10%, 5%, and 1% of the original sequencing depth. For each downsampled dataset, we applied the same pre-processing, PCA, and clustering steps as the original RA dataset. We used the PCA results for each downsampled dataset and the original RA dataset’s clustering assignments as input for **Step 2**.

For dataset generation, we used the same study design as the baseline analyses. We ran 500 simulations for each combination of parameters.

### 4.11 Analyses containing fewer batches (increased samples per batch)

For these analyses, we used almost the same parameters as the baseline analyses. However, we varied the number of samples that were placed into each batch, so that the studies had 2 (50 samples per batch), 5 (20 samples per batch), 10, 15, or 25 batches overall. We also varied the values of b_scale_: 0, 0.5, 1, 2, and 4. We ran 500 simulations for each combination of parameters.

### 4.12 Multiplexing analyses

For the sequential study design, we assigned 100 samples (2000 cells each) into 100 batches using the provided “distribSamplePerBatch” function so that each batch contained cells from only 1 sample (2000 cells per batch). We induced fold-changes into a randomly chosen cluster, and we set b_scale_, s_scale_, and cf_scale_ all equal to 1.

For the multiplexing study design, we split 100 samples (2000 cells each) into 4 equally sized subsamples (500 cells each). We then assigned these subsamples into 100 batches using the provided “distribSplitSample” function so that each batch contained cells from 4 different subsamples (2000 cells per batch). We induced fold-changes into a randomly chosen cluster, and we set b_scale_, s_scale_, and cf_scale_ all equal to 1. We ran 500 simulations for each combination of parameters for both analyses.

### 4.13 Increasing the total number of samples versus number of cells per sample

For these analyses, we used the same study design as the baseline analyses, but we varied the total number of samples (10, 20, 40, 80, or 160) and the number of cells per sample (25, 50, 100, 200, 400, 800, and 1600). For the main figures, we show results for fold-changes of 1.5 and 2 for the RA and TB settings, and results for fold-changes of 2 and 4 for the UC setting. We provide power curves for the rest of the fold-changes in supplement. We ran 100 simulations for each combination of parameters.

## Data Availability

Processed scRNA-seq data for the RA dataset is available at https://www.immport.org/shared/study/SDY998, with raw scRNA-seq deposited at dbGaP https://www.ncbi.nlm.nih.gov/projects/gap/cgi-bin/study.cgi?study_id=phs001457.v1.p1. Processed scRNA-seq data for the UC dataset is available at Single Cell Portal: SCP259. We provide processed meta-data tables (including cell state cluster assignments) and gene expression matrices for the RA and UC datasets at https://github.com/immunogenomics/scpost. The memory T cell CITE-seq data from the TB dataset is currently private and will be made available at GEO accession GSE158769.

## Code Availability

An R implementation of the scPOST framework (along with input prototype data structures from the RA and UC datasets) is provided at https://github.com/immunogenomics/scpost.

## Acknowledgements

This work is supported in part by funding from the National Institutes of Health (UH2AR067677, 1U01HG009088, U01 HG009379, and 1R01AR063759). We thank members of the Raychaudhuri Lab. This work was supported by the Accelerating Medicines Partnership (AMP) in Rheumatoid Arthritis and SLE Network. AMP is a public-private partnership (AbbVie Inc., Arthritis Foundation, Bristol-Myers Squibb Company, Janssen Pharmaceuticals, Lupus Foundation of America, Lupus Research Alliance, Merck Sharp & Dohme Corp., National Institute of Allergy and Infectious Diseases, National Institute of Arthritis and Musculoskeletal and Skin Diseases, Pfizer Inc., Rheumatology Research Foundation, and Sanofi and Takeda Pharmaceuticals International, Inc.) created to develop new ways of identifying and validating promising biological targets for diagnostics and drug development. Funding was provided through grants from the National Institutes of Health (UH2-AR067676, UH2-AR067677, UH2-AR067679, UH2-AR067681, UH2-AR067685, UH2-AR067688, UH2-AR067689, UH2-AR067690, UH2-AR067691, UH2-AR067694, and UM2-AR067678). We thank mambers of the Tuberculosis Research Unit (TBRU) LIMAA and Socios En Salud, in particular D. Branch Moody, Megan Murray, Jessica Beynor, Yuriy Baglaenko, Sara Suliman, Ildiko van Rhijn, and Leonid Lecca, for their contributions to generating the memory T cell dataset. The content is solely the responsibility of the authors and does not necessarily represent the official views of the National Institutes of Health.

## Author Contributions

N.M., C.Y.F., and S.R. conceived the project. N.M. and I.K. developed the method and performed the analyses under the guidance of S.R. All authors participated in interpretation and writing of the manuscript.

## Competing Interests

S.R. receives research support from Biogen

## Notes

### Competing Interest Statement

The authors have declared no competing interest.

