## Supplemental Figures 1-11 for "Maximizing statistical power to detect clinically associated cell states with scPOST"

### Supplementary Figures and Tables

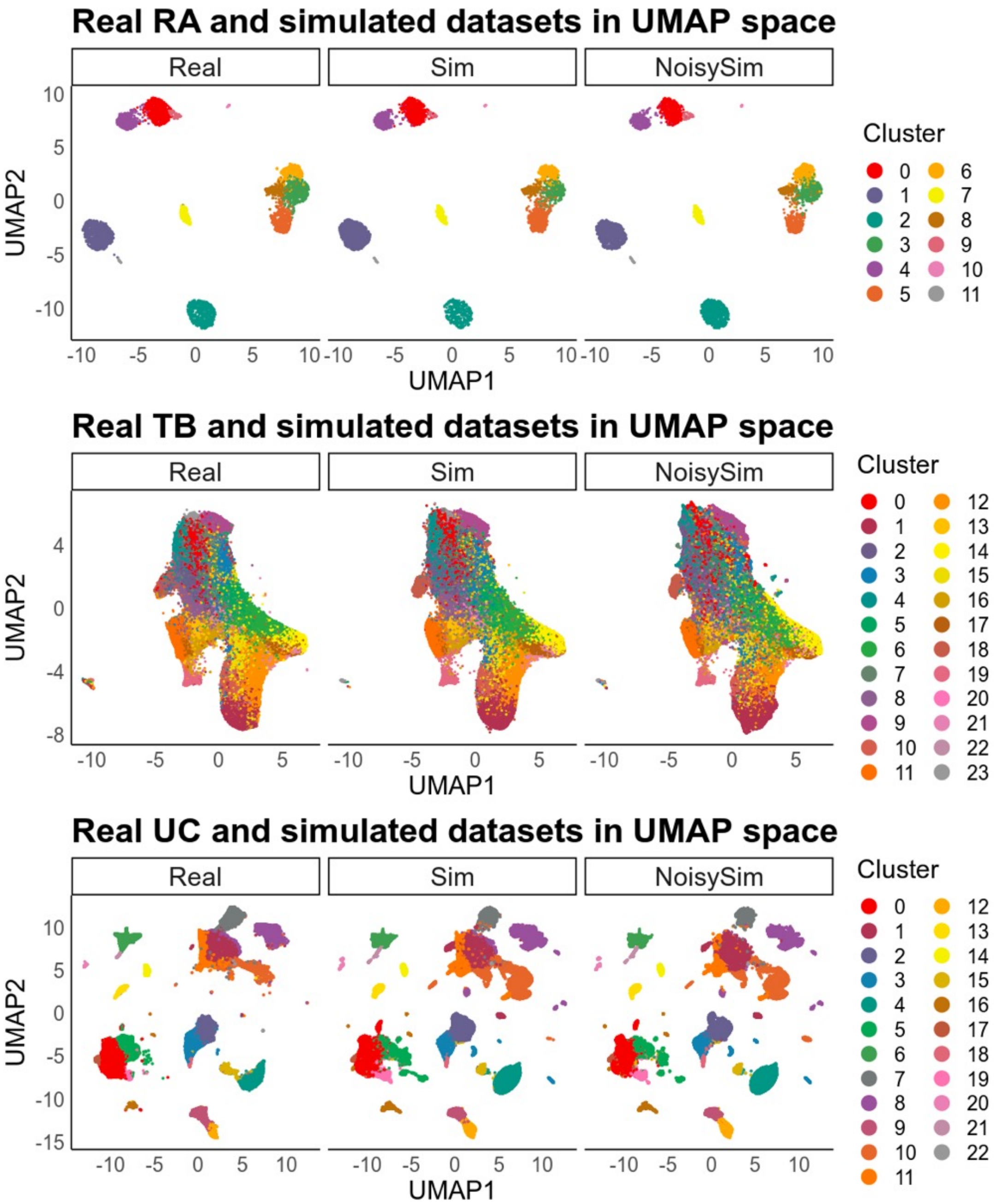


**Supplementary Figure 1 | scPOST can generate single-cell datasets similar to the original input prototype dataset, as well as datasets with different magnitudes of noise.** UMAP visualizations of the RA, TB, and UC datasets. For each dataset, each panel is embedded in the same UMAP space. The “Real” panel contains cells from the original input prototype dataset, the “Sim” panel contains cells from a simulated realistic dataset, and the “NoisySim” panel contains cells from a simulated dataset that contained 4x the estimated batch-associated variance ($b_{scale}$ = 4).

| | RA dataset | TB dataset | UC dataset |
|---|---|---|---|
| Single-cell technology | CelSeq2[41] | 10X Chromium 3' v3 | 10X Chromium 3' v2/v3 |
| Number of samples | 21 | 259 | 30 |
| Number of cells passed QC | 5,265 | 496,517 | 235,229 |
| Type of samples | Synovial tissue (joint replacement procedure or biopsy) | PBMCs (blood) | Intestinal biopsy |
| Broad cell types assayed | Immune and stromal cells (T/B cells, Monocytes, Fibroblasts) | Memory T cells | Immune and stromal cells (T/B/Myeloid cells, Fibroblasts, Endothelial cells, Epithelial cells) |
| Mean reads/cell | 7,300 | 4,920 | 4,582 |
| Mean unique genes/cell | 2,432 | 1,472 | 988 |
| Batch-correction | Harmony | None | None |

**Supplementary Table 1 | Characteristics of the RA, TB, and UC datasets.**

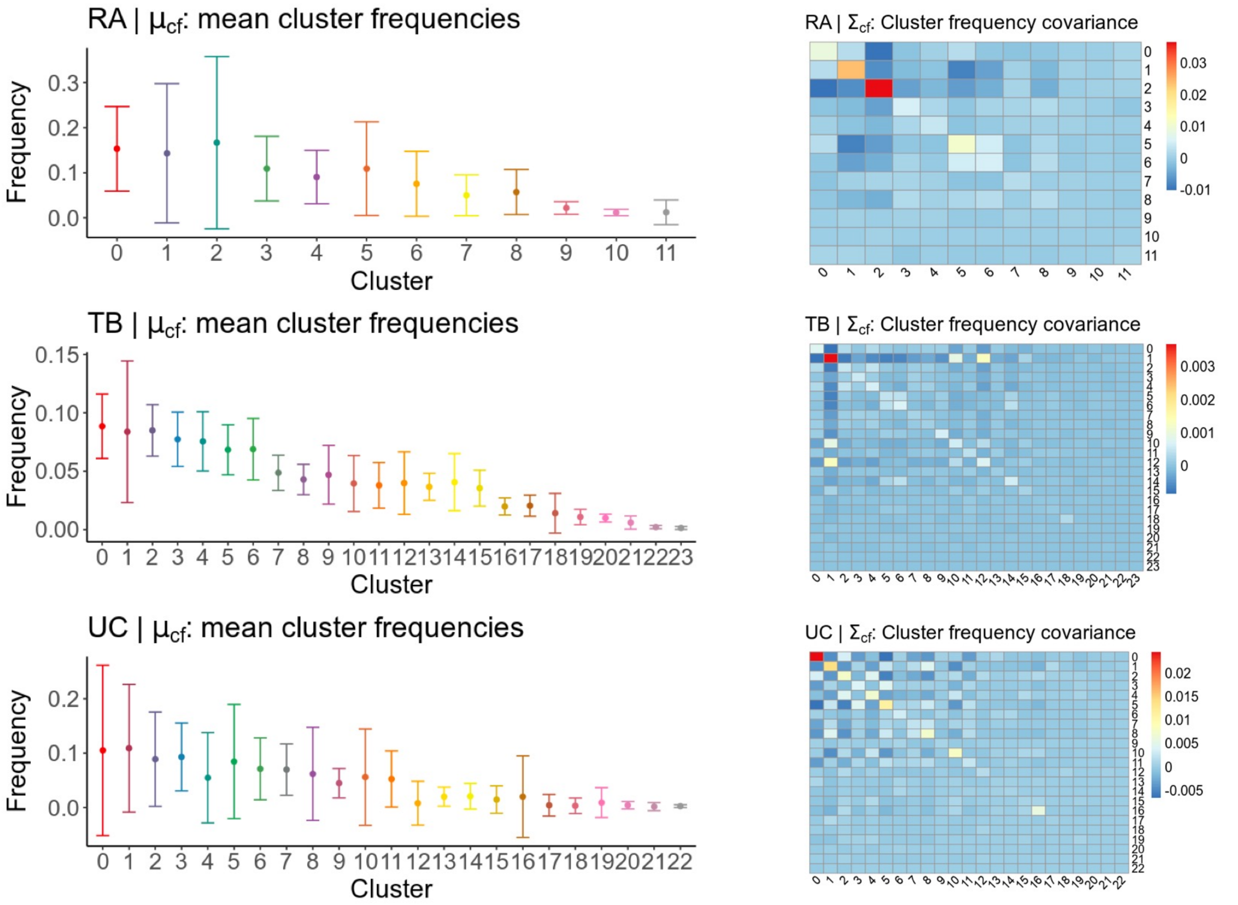


**Supplementary Figure 2 | Cell state frequency variation across samples in each of dataset differences between datasets. Left:** Frequency distribution for each cell state cluster in the RA, TB, and UC datasets. The plotted mean frequencies are the observed mean frequency of that cluster across all samples in their respective dataset. In all panels, error bars represent one standard deviation from the mean in each direction. **Right:** Cell state cluster frequency covariance matrices for the RA, TB, and UC datasets.

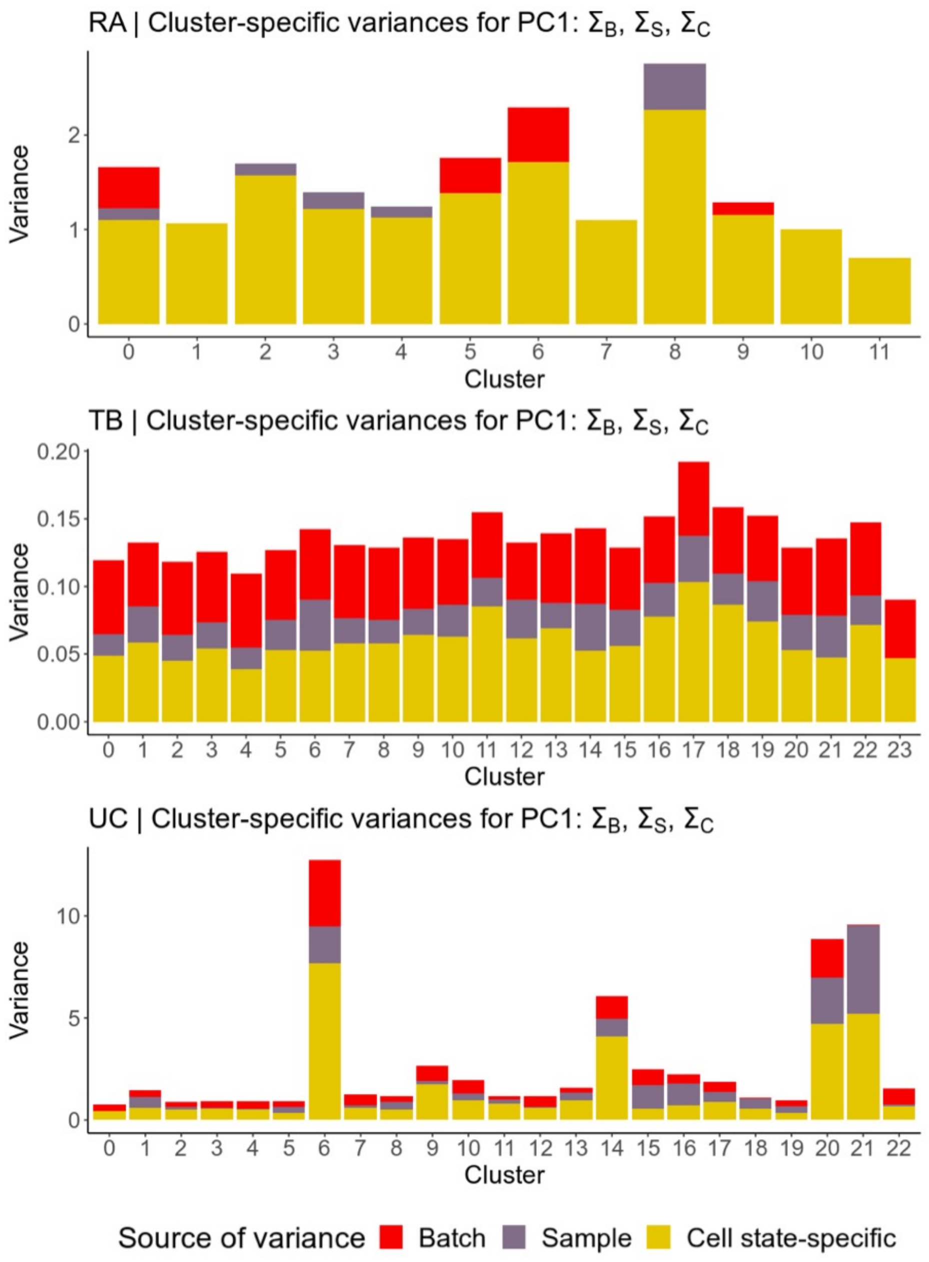


**Supplementary Figure 3 | Estimated parameters from the input prototype RA, TB, and UC datasets. A-C,** Bar plots of the estimated variance in each cell state cluster for the first principal component in each of the RA, TB, and UC datasets. Each bar is colored by the source of noise contributing to that variance.

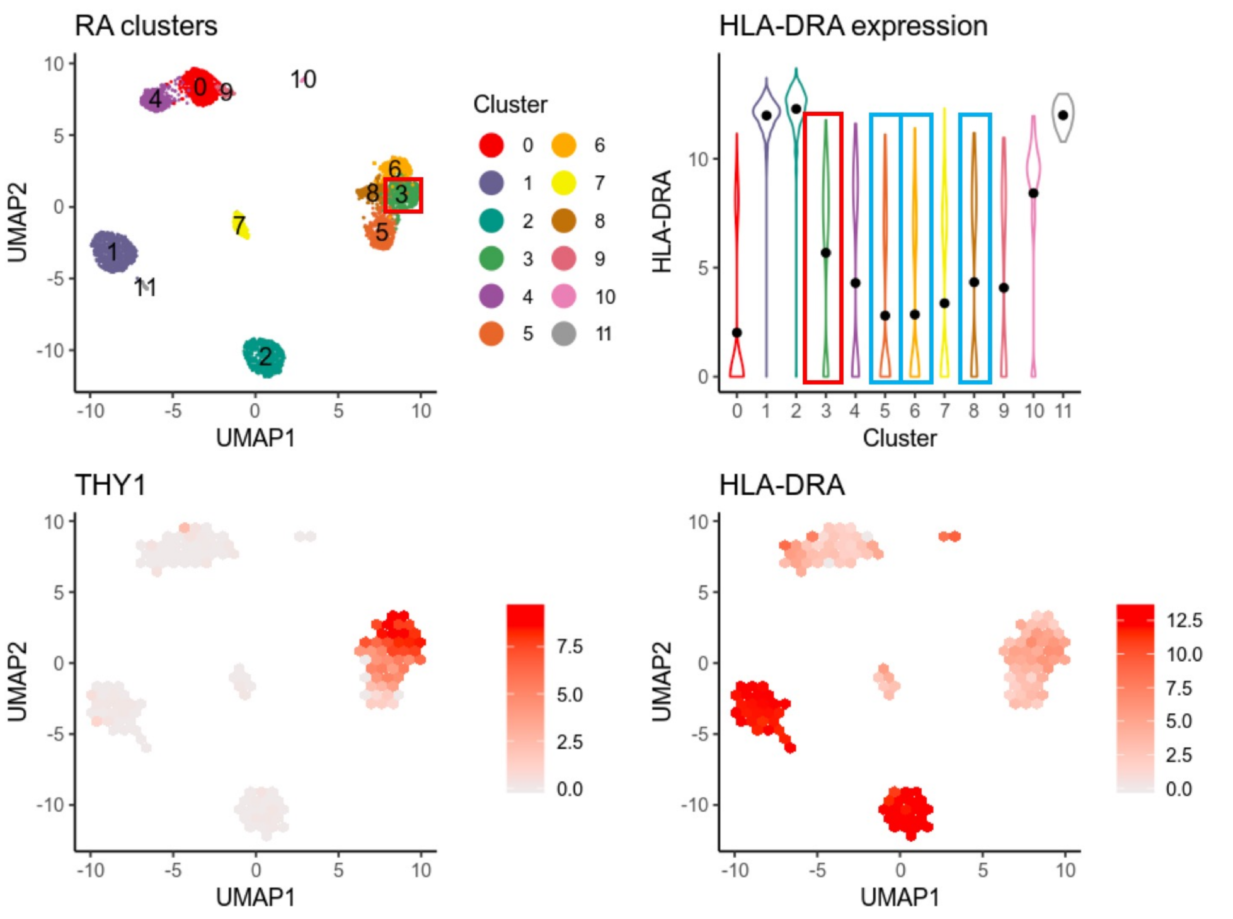


**Supplementary Figure 4 | The RA dataset contains THY1+ sub-lining fibroblasts that express HLA-DRA.** For our analyses in which we expanded the RA study, we focused on the THY1+ sub-lining fibroblast cluster (cluster 3 highlighted in red, other fibroblast clusters highlighted in blue) that most highly expressed *HLA-DRA* that corresponds to the HLA-DRA$^{hi}$ fibroblast population described in the paper. The RA dataset was filtered to only include fibroblasts before inputting into scPOST for Fig. 4a.

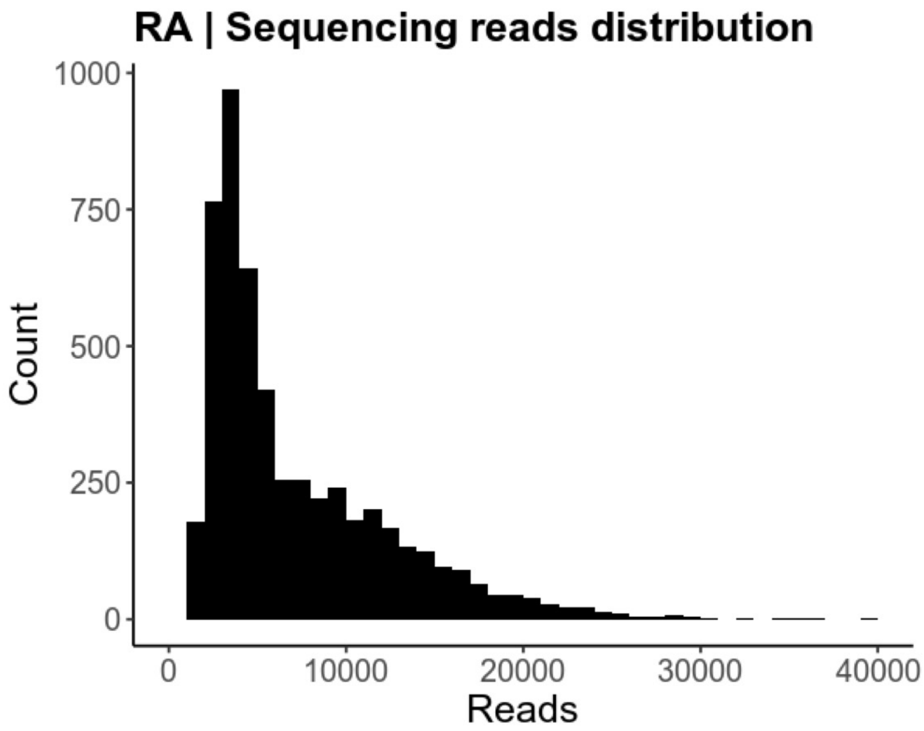


**Supplementary Figure 5 | Sequencing read distribution for the RA dataset.** Histogram of the sequencing read distribution of the original RA dataset. Downsampled datasets maintained similar sequencing read distributions, but with a lower mean number of reads.

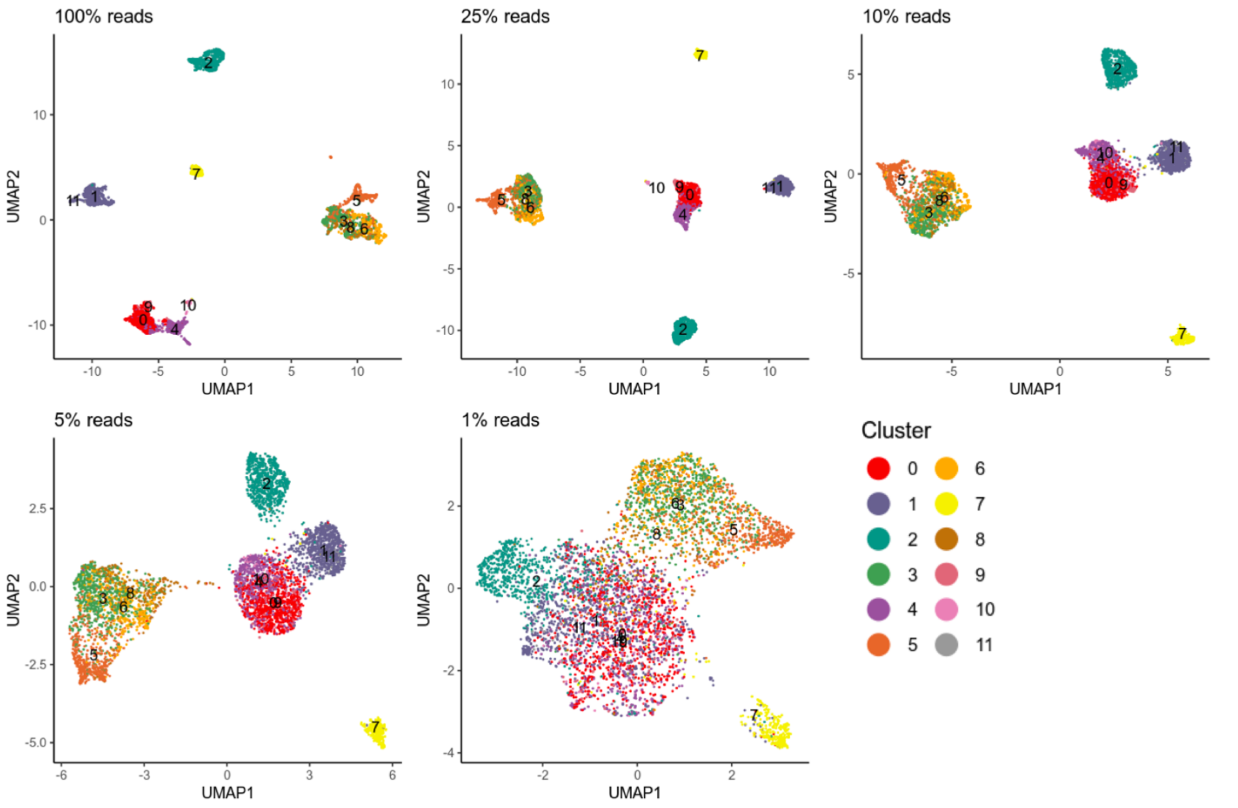


**Supplementary Figure 6 | UMAP visualizations of datasets derived from downsampling the RA dataset's gene expression read data.** The RA dataset's sequencing reads were downsampled to have a specific percentage of reads for each count. Downsampled datasets were input into standard PCA dimensionality reduction pipeline, and then visualized with UMAP. Cells are colored by the cell state identities obtained from the original RA dataset.

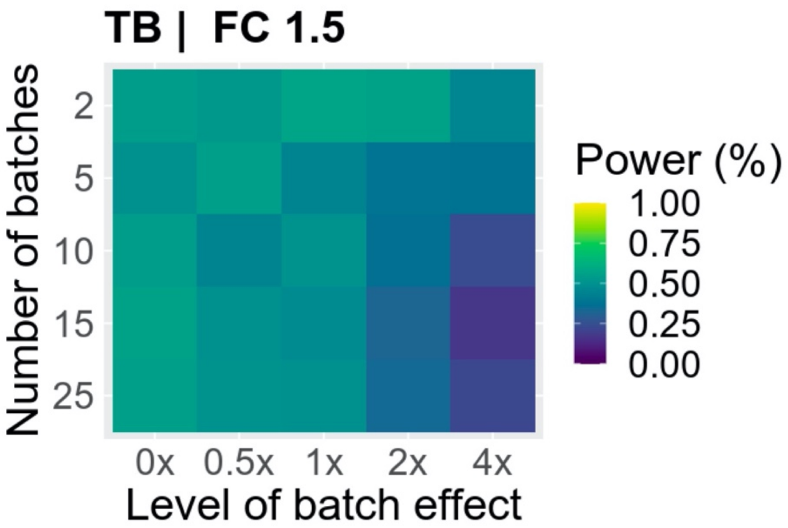


**Supplementary Figure 7 | Decreasing the number of batches (by increasing the number of samples per batch) provides negligible benefit until high magnitude of batch effects.** Power calculations across different ranges of scaled batch effects and number of batches, with the induced fold change set to 1.5. Simulations were performed in the realistic context, but with modulated levels of batch-influenced variation on gene expression. The number of batches decreasing indicates increasing the number of samples run in a batch.

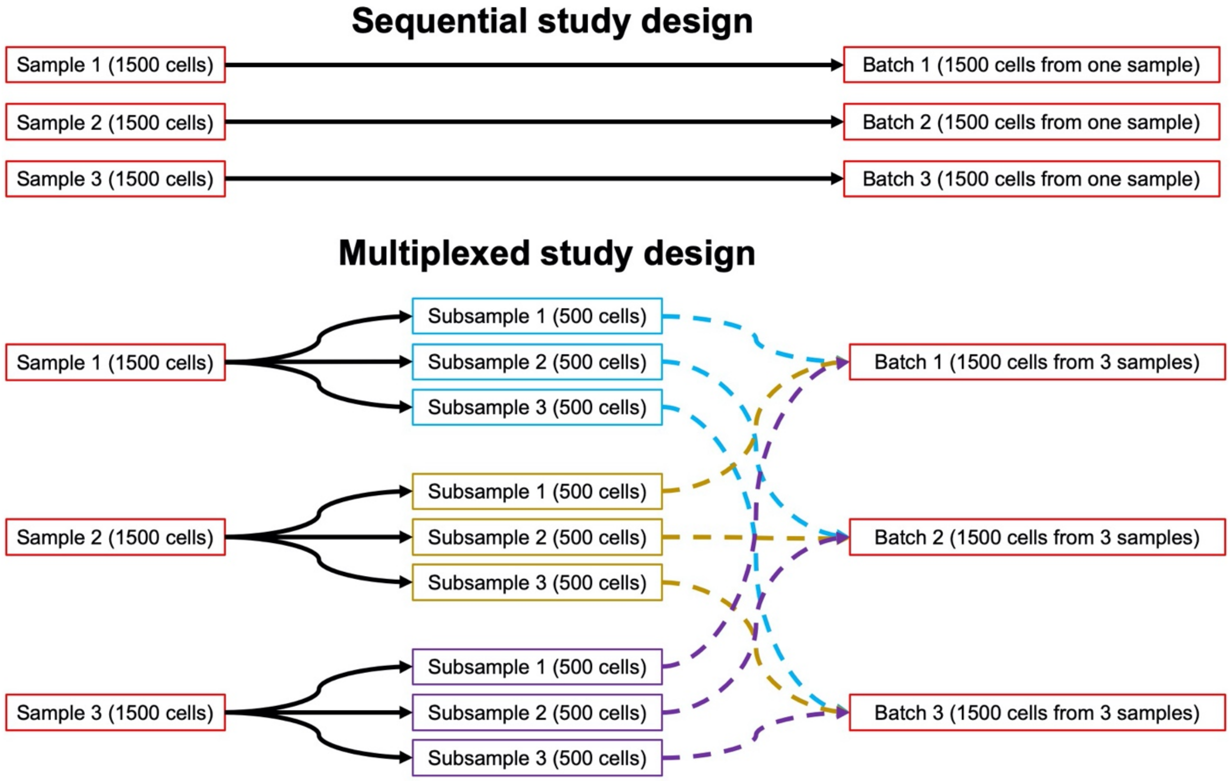


**Supplementary Figure 8 | scPOST facilitates exploration of different study designs, such as batch multiplexing structures.** In Fig. 5A, we compare the estimated power from two batch multiplexing structures with scPOST. The non-multiplexed sequential design placed each simulated sample in its own batch, so that each batch contained cells from only one sample. In Fig. 5a, we placed samples (total of 100 with 2000 cells each) into 100 batches. The multiplexed study design featured a batch structure in which each simulated sample was split into equally-sized subsamples. These subsamples were then placed into different batches, so that each batch contained cells from multiple samples. In Fig. 5a, we split each sample (total of 100 with 2000 cells each) into 4 subsamples (500 cells each), which were then placed into 100 batches (each batch contained cells from different subsamples).

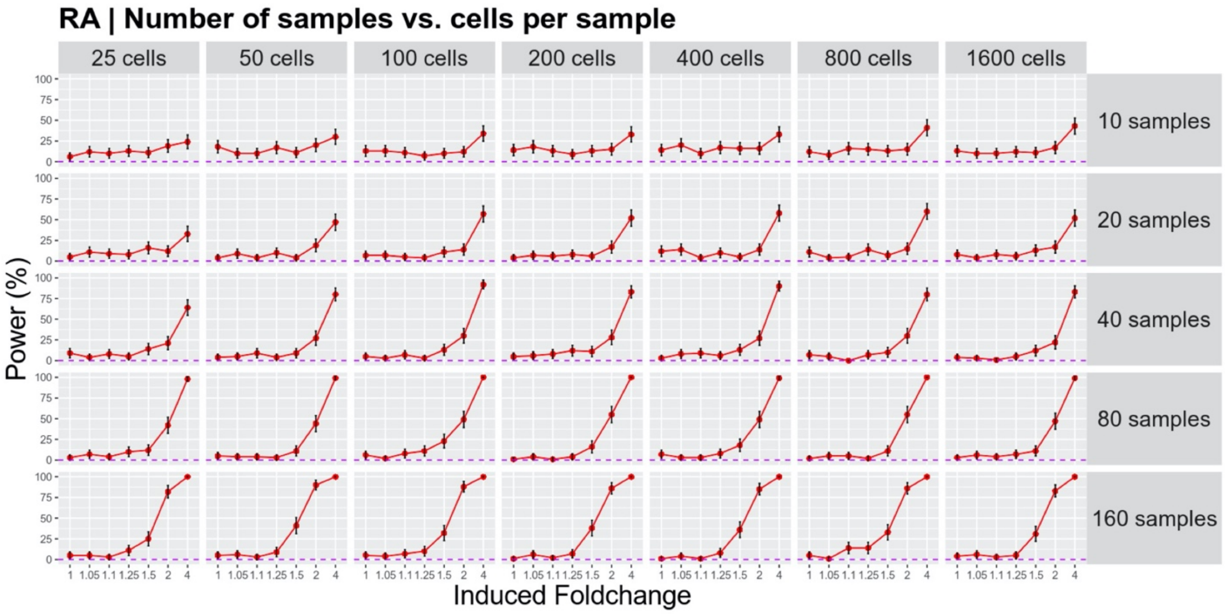


**Supplementary Figure 9 | Power estimations in the RA setting for dataset sizes characterized by a variable number of samples and cells per sample.** Power estimations for different sample/cell per sample combinations. Simulations were performed in the baseline context (Fig. 4b) with each data point representing 100 simulations. Grid elements along a top right-bottom left diagonal represent an equivalent number of cells. Error bars represent 95% binomial proportion confidence intervals and the dotted horizontal purple line represents 5% power.

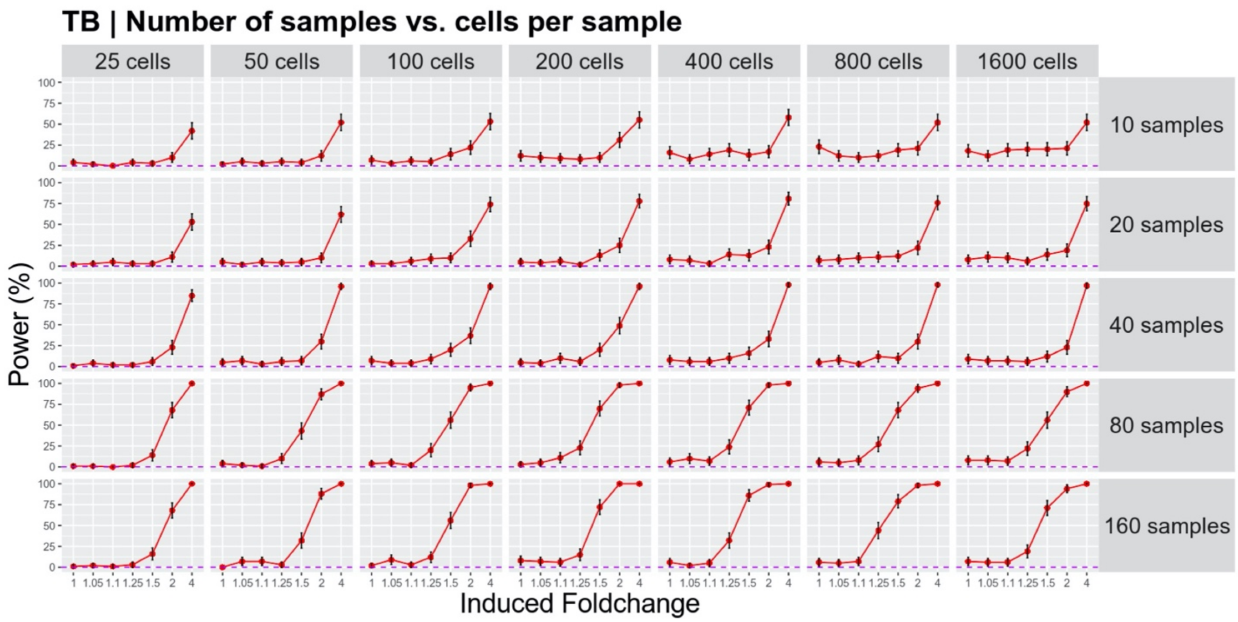


**Supplementary Figure 10 | Power estimations in the TB setting for dataset sizes characterized by a variable number of samples and cells per sample.** Power estimations for different sample/cell per sample combinations. Simulations were performed in the baseline context (Fig. 4c) with each data point representing 100 simulations. Grid elements along a top right-bottom left diagonal represent an equivalent number of cells. Error bars represent 95% binomial proportion confidence intervals and the dotted horizontal purple line represents 5% power.

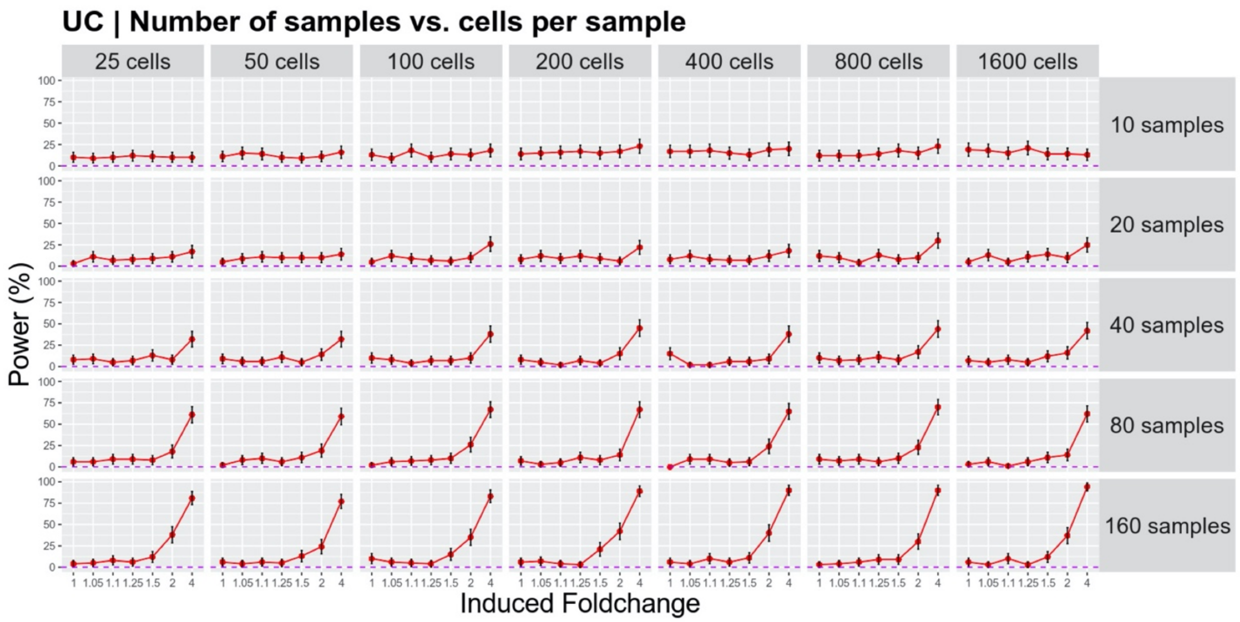


**Supplementary Figure 11 | Power estimations in the UC setting for dataset sizes characterized by a variable number of samples and cells per sample.** Power estimations for different sample/cell per sample combinations. Simulations were performed in the baseline context (Fig. 4d) with each data point representing 100 simulations. Grid elements along a top right-bottom left diagonal represent an equivalent number of cells. Error bars represent 95% binomial proportion confidence intervals and the dotted horizontal purple line represents 5% power.
